# Effects of Social Organization, Trap Arrangement and Density, Sampling Scale, and Population Density on Bias in Population Size Estimation Using Some Common Mark-Recapture Estimators

**DOI:** 10.1101/114306

**Authors:** Manan Gupta, Amitabh Joshi, T. N. C. Vidya

## Abstract

Mark-recapture estimators are commonly used for population size estimation, and typically yield unbiased estimates for most solitary species with low to moderate home range sizes. However, these methods assume independence of captures among individuals, an assumption that is clearly violated in social species that show fission-fusion dynamics, such as the Asian elephant. In the specific case of Asian elephants, doubts have been raised about the accuracy of population size estimates. More importantly, the potential problem for the use of mark-recapture methods posed by social organization in general has not been systematically addressed. We developed an individual-based simulation framework to systematically examine the potential effects of type of social organization, as well as other factors such as trap density and arrangement, spatial scale of sampling, and population density, on bias in population sizes estimated by POPAN, Robust Design, and Robust Design with detection heterogeneity. In the present study, we ran simulations with biological, demographic and ecological parameters relevant to Asian elephant populations, but the simulation framework is easily extended to address questions relevant to other social species. We collected capture history data from the simulations, and used those data to test for bias in population size estimation. Social organization significantly affected bias in most analyses, but the effect sizes were variable, depending on other factors. Social organization tended to introduce large bias when trap arrangement was uniform and sampling effort was low. POPAN clearly outperformed the two Robust Design models we tested, yielding close to zero bias if traps were arranged at random in the study area, and when population density and trap density were not too low. Social organization did not have a major effect on bias for these parameter combinations at which POPAN gave more or less unbiased population size estimates. Therefore, the effect of social organization on bias in population estimation could be removed by using POPAN with specific parameter combinations, to obtain population size estimates in a social species.

## Introduction

Population size estimation is a vital aspect of studying the ecology of animal populations in the wild. It is needed to study population dynamics [1,2], to estimate the sex-ratio of populations [3,4], to monitor populations facing the risk of extinction [5], and to also calculate effective population size for studying evolutionary dynamics or ascertaining the threat to genetic diversity due to drift [6,7]. The standard practice for estimating population size is to use either direct counts, transect counts, or mark-recapture methods [8]. Direct counts only work in the case of easily detectable species in which detection of all individuals in the study area is possible. Transect counts give reliable results only when the study population is closed, implying no addition or removal of individuals due to demographic (birth and death) or dispersal (immigration and emigration) events [8]. Mark-recapture methods can take into account detection probabilities less than one, as well as the non-closure (openness) of populations with regard to demography and dispersal [8,9]. The above advantages make mark-recapture methods a viable option to use for estimating population sizes of many species. However, the use of mark-recapture methods in the case of social species remains controversial [8]. The problem is that mark-recapture methods typically assume the detection probabilities of individuals to be independent [8–10], whereas this assumption is likely to be violated in social species that show coordinated movement of socially interacting individuals. The effects of such non-independence of capture probabilities can be non-trivial: Boulanger *et al.* [11] found that non-independence between pairs of individuals (mother-offspring pairs in their case) caused population size estimates to be biased, despite incorporating heterogeneity in capture probabilities between those individuals in their simulation. However, mark-recapture methods can still be applied to social species with fixed groups by assuming the detection probabilities to be independent among groups. Then, the capture of entire groups rather than individuals is considered, and the estimate of total groups can be multiplied by the mean group size to get an estimate of total population size [8]. In cases where this approach cannot be used, corrections need to be made for the variance of mark-recapture parameters. Non-independence of detection causes over-dispersion in multinomial data [12], resulting in smaller than actual variance for estimates of mark-recapture parameters [13]. To obtain unbiased estimates of the variance in such situations, the variance needs to be inflated by multiplying it by a variance inflation factor (*ĉ*), which is derived from the goodness-of-fit statistic [14]. This method, too, does not always yield a proper estimate of *ĉ*, especially when sample sizes are relatively small [15]. Consequently, more sophisticated numerical methods are called for in such cases [13,15].

The problems discussed above make it especially difficult to obtain robust estimates of population sizes for species showing fission-fusion dynamics, in which groups of individuals may come together or split away, changing spatio-temporal cohesiveness depending on resources and competition [16–18]. Female Asian elephants (*Elephas maximus*) are a case in point. Asian elephants live in female-bonded groups, with adult males leading largely solitary lives [19–23]. Consequently, the movement of individual females is not independent, as females in a group tend to move together. The smallest unit of social organisation in female Asian elephants is the mother-offspring unit, several of which may join together to form a family group [19,20,22,23]. These family groups or their subsets may further associate to form higher levels of organisation referred to as ‘bond groups’ or ‘clans’ [20]. Asian elephant societies show fission-fusion dynamics [19,20,24,25] and, consequently, group identities are not conserved over time. This also rules out the possibility of carrying out mark-recapture analysis on fixed groups. Most studies that have estimated population size for Asian elephants have not used mark-recapture methods [26–29]. In some studies, population sizes of only adult males were estimated by photographic mark-recapture [30] or sight-resight mark-recapture [31]. In some other studies, for want of better methods, population size of adult females or the total population size were estimated by mark-recapture methods despite the non-independence of detection probabilities among individual females [32–35]. It is important to assess the reliability of using mark-recapture methods in this endangered species, whose global estimates are presently thought to be only educated guesses [36,37], so that monitoring can be carried out based on sound methods.

Here, we report the development of an individual-based simulation of animal movement, drawing on empirical information from field studies of the Asian elephant in the Nagarahole and Bandipur National Parks and other parts of the Nilgiris-Eastern Ghats Reserve in southern India. We used this simulation framework to test whether mark-recapture models give robust and unbiased estimates of population size in the case of social species in which individuals show coordinated movement to varying degrees. We specifically tested for bias in the commonly used POPAN estimator [38] and Robust Design estimators with and without detection heterogeneity [39–43], using data obtained from simulations with varying social organizations, population densities, trap densities, spatial trap arrangements, and spatial scales of sampling. We considered all combinations of two trap arrangements (uniform and random), four trap densities (0.1, 0.5, 1.0, and 1.5 traps Km^−2^), three sampling scales (dividing up a fixed study area into 4 large, 9 medium, or 16 small sampling blocks) and three social organizations (individuals only, individuals within clans, and individuals within groups within clans). In all simulations, only adult females were considered and henceforth we refer to adult female density as adult density (see Methods for details and rationale). For each combination of these factor-levels, adult density was treated as a classifying factor in two alternate ways. In one set of analyses, we used the initial adult density assigned to the entire fixed study area considered in the simulation. In a second set of analyses, we used the actual mean adult density observed in each sampling block in the simulation, averaged over time (i.e. different sampling points: see Methods for details and rationale). Thus, we carried out two complete sets of analyses, using either initial adult density or actual adult density, respectively. We report here, the results only from the analyses performed using actual density as a factor, as the results based on actual and initial density were very similar (see Methods for details and rationale, S1 Appendix for results based on initial density). We found that social organization significantly affected bias in population estimation under most cases, although the magnitude of bias depended on other parameter combinations. POPAN clearly outperformed the two Robust Design methods (with or without detection heterogeneity) we used, yielding far more unbiased estimates of population size. Bias from social organization could be avoided by using POPAN with random trap arrangement and relatively high trap densities. While we carried out this study with the specific aim of assessing commonly used mark-recapture methods in the context of population size estimation for Asian elephant populations in the wild, we expect that the simulation framework we have developed will be applicable to many other species of medium to large sized mammals with relatively minor modifications. The simulation framework could also be expanded upon to address a number of questions pertaining to sampling effects in the context of different data sampling approaches used in observational studies of wild mammals that have different patterns of coordinated movement in response to varying resource distributions.

## Results

### Observed AI Distribution

We used different patterns of Association Probabilities (APs; see Methods) to determine how individuals of the same group, of different groups but within the same clan, or of different clans tended to move together. In order to check whether these APs actually yielded characteristic signatures of the three social organizations we tried to simulate, we examined Association Indices (AIs; see Methods) for the three social organizations over two different time periods in the simulations (Fig 1). AIs among individuals of the same group or the same clan were almost always zero for the non-associating individuals case, implying that, as expected, individuals exhibiting a lack of social organization rarely ever moved together. The groups within clans social organization was meant to reflect a pattern of coordinated movement wherein group-mates would spend a lot of time close to each other, whereas clan-mates from different groups would encounter each other only occasionally. The AI distribution within clans for this social organization was seen to be relatively flat, with relatively greater frequencies around zero and one, respectively, during both time periods examined (Fig 1). This observation is concordant with the expectation that group-mates within clans should show relatively high AIs, whereas clan-mates from different groups should only rarely associate with one another. The fixed clans social organization was meant to simulate a situation wherein, irrespective of group, clan-mates would tend to move together with one another much more than they would with members of another clan. For this pattern of APs, within-clan AIs were mostly between 0.9 and 1 (Fig 1), thus reflecting the desired type of social organization. Some pairs, however, showed very low within-clan AI values for the fixed clans case (Fig 1), which is likely due to the *K*-means algorithm disintegrating bigger clans into multiple clusters, and also due to the relatively rare event when an individual started to move away from the clan. In case of the within-group AI distributions, both the groups within clans and fixed clans social organizations showed distributions with many high values, and a few low ones, similar to one another and to the AI distribution within clans for the fixed clans case (Fig 1). This is concordant with the fact that the clans in the fixed clans case were intended to just be larger versions of the groups in the groups within clans case. On the whole, the AI distribution data indicate that the different patterns of APs we used in our simulations did, in fact, result in AI distributions reflecting the three social organizations intended.

**Fig 1.**
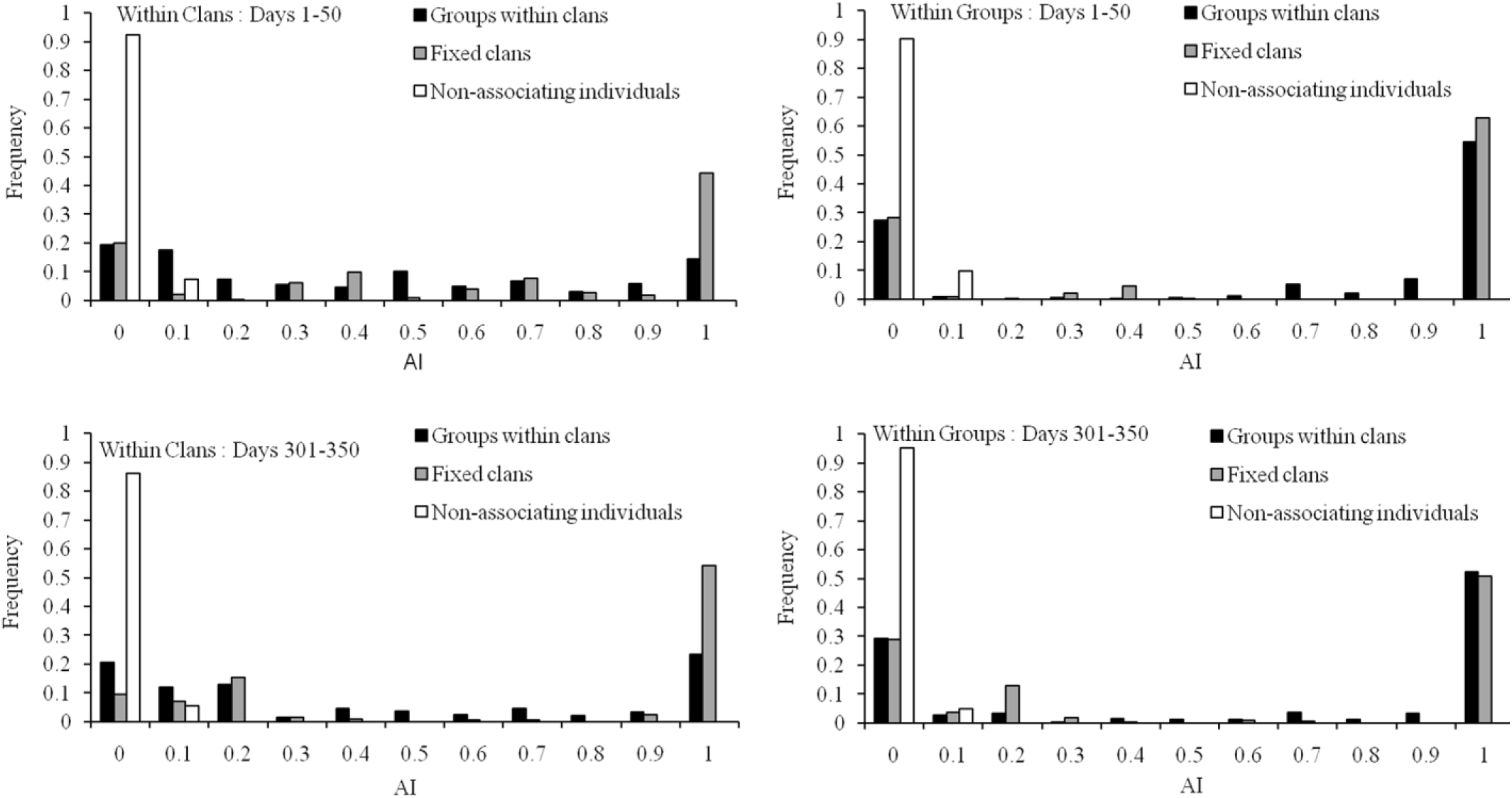
Frequency distributions of calculated association indices (AIs) within clans and within groups for the three social organizations during the first and last 50 days of the 350 day period over which AI patterns were assessed.

### Bias in Population Size Estimation

Overall, population size estimates from all three mark-recapture models used tended to be underestimates. The relative bias values were, by and large, negative for all factor-level combinations, regardless of which mark-recapture model was used (Figs 2–4: relative bias values ranged from about −0.99 to +0.06, with most values being negative; results are only shown for actual average adult density in each sampling block, see S1 Appendix for results with initial adult density as a factor). Using POPAN (Fig 2) yielded smaller absolute levels (ignoring the direction of bias) of relative bias values than either Robust Design (Fig 3) or Robust Design with Heterogeneity (Fig 4), with the latter two models always yielding relative bias values more negative than −0.40. Overall, the least biased estimates of population size (relative bias close to zero) were obtained using POPAN with a random trap arrangement, high trap density, and high actual density (Fig 2). In the case of all three mark-recapture models, five-way analyses of variance (ANOVAs) revealed that trap arrangement (uniform or random) explained the majority of variation in relative bias (SI Appendix: Table M for POPAN; Table O for Robust Design; Table Q for Robust Design with Heterogeneity). Consequently, four-way fully-factorial ANOVAs were also performed within each trap arrangement factor level (uniform or random) in order to better tease apart the effects of factors other than trap arrangement. In this section, we have presented the data on mean relative bias in population size estimation graphically (Figs 2–4) and summarized the important results from the five-way and four-way ANOVAs (Table 1). S1 Appendix contains the complete set of ANOVA tables for all four- and five-way ANOVAs (Tables M-AD in S1 Appendix), as well the means for various combinations of factor levels in two- and three-way interactions, along with the results of pair-wise comparisons among them (Tables A-L in S1 Appendix). The results of the various analyses for each mark-recapture model are separately discussed below.

**Table 1.** The pattern of significance in the four-way fully-factorial ANOVAs done on relative bias in population size estimates using the three mark-recapture models using actual density as a factor (***=*P*<0.001, **=*P*<0.01, *=*P*<0.05, Blank cell=Not significant). Effect sizes (*η^2^* = _SSFactor or Interaction_/SS_Total_) are shown as colour shading. TA: trap arrangement.

|  | POPAN |  | Robust Design |  | Robust Design with Heterogeneity |  |
| --- | --- | --- | --- | --- | --- | --- |
|  | Uniform TA | Random TA | Uniform TA | Random TA | Uniform TA | Random TA |
| **Trap Density (1)** | *** | *** | *** | *** | *** | *** |
| **Spatial Scale (2)** | *** |  | *** | *** | *** |  |
| **Social org. (3)** | *** | *** | *** | *** | *** | *** |
| **Adult Density (4)** | *** | *** | *** | *** | *** | *** |
| **1x2** |  | *** | *** | *** | *** | *** |
| **1x3** |  | *** | *** | *** | *** | *** |
| **2x3** | *** | *** |  | *** |  | *** |
| **1x4** | *** | *** | *** | *** | *** | *** |
| **2x4** | *** | * | *** | *** | *** |  |
| **3x4** | *** | *** | *** | *** | *** |  |
| **1x2x3** |  |  |  | *** | ** | *** |
| **1x2x4** |  | ** | *** | *** | *** |  |
| **1x3x4** |  |  |  | *** | * | *** |
| **2x3x4** |  | *** | * | *** |  | *** |
| **1x2x3x4** |  |  | * |  |  | * |

**Fig 2.**
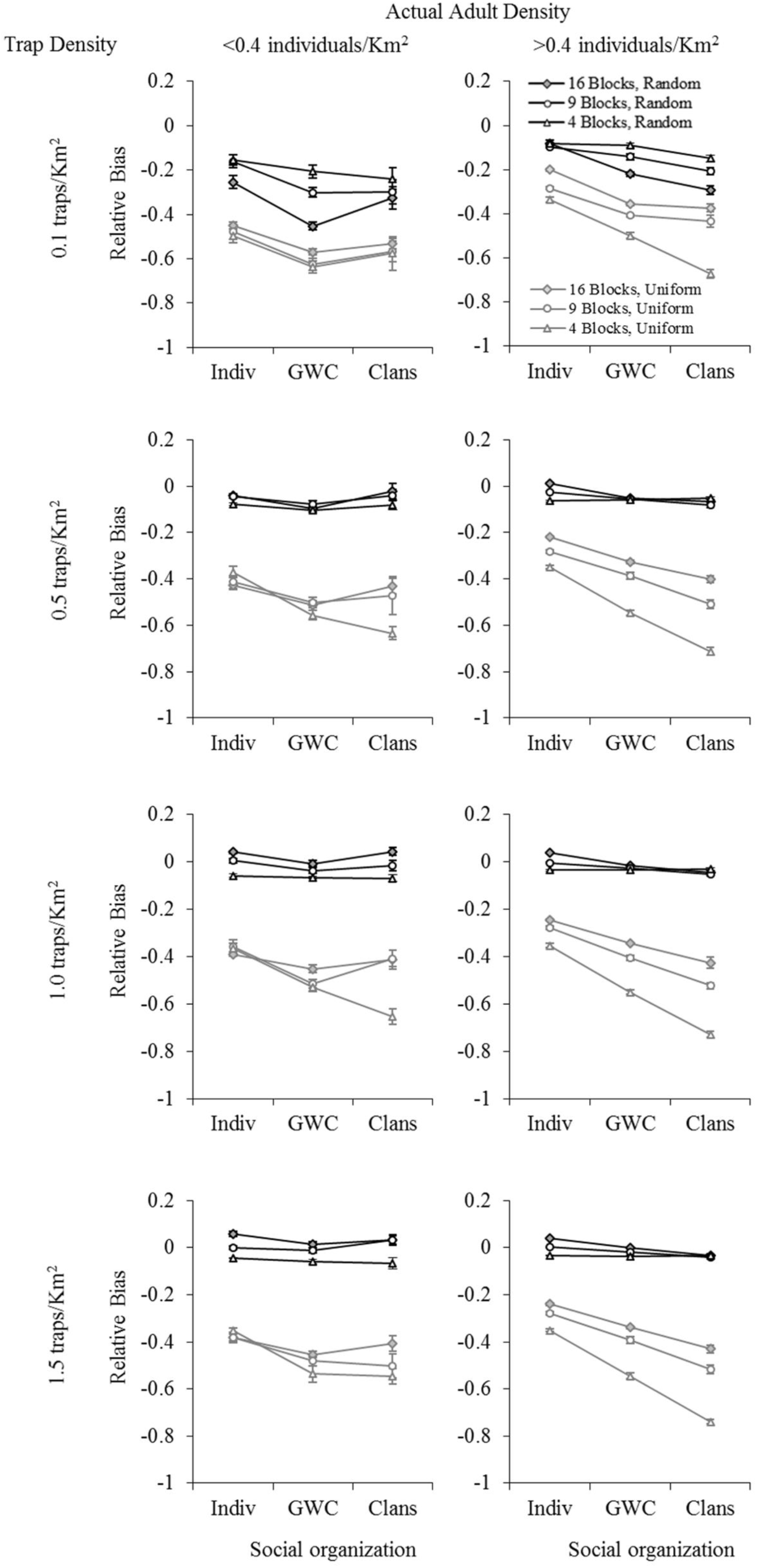
The effect of trap density, sampling scale, social organization, and adult density on mean relative bias in population size estimation using POPAN. Data shown are for a uniform (UTA, grey lines) and random (RTA, black lines) trap arrangement, using actual adult densities. Error bars represent standard errors. B = Blocks (i.e., 16 Blocks, etc.). Social organization: Indiv: non-associating individuals; GWC: groups within clans; Clans: fixed clans.

**Fig 3.**
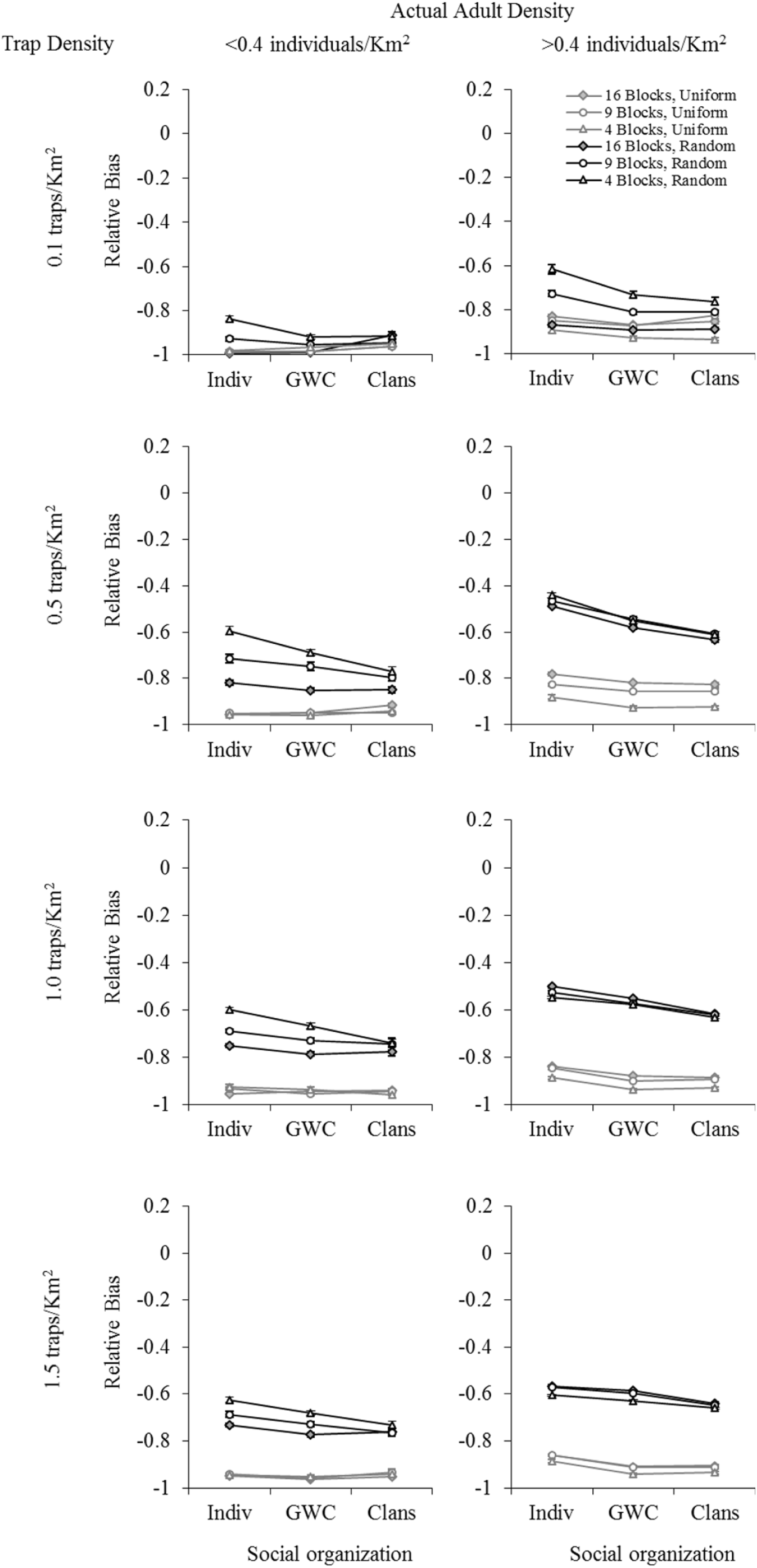
The effect of trap density, sampling scale, social organization, and adult density on mean relative bias in population size estimation using Robust Design. Data shown are for a uniform (UTA, grey lines) and random (RTA, black lines) trap arrangement, using actual adult densities. Error bars represent standard errors. B = Blocks (i.e., 16 Blocks, etc.). Social organization: Indiv: non-associating individuals; GWC: groups within clans; Clans: fixed clans.

**Fig 4.**
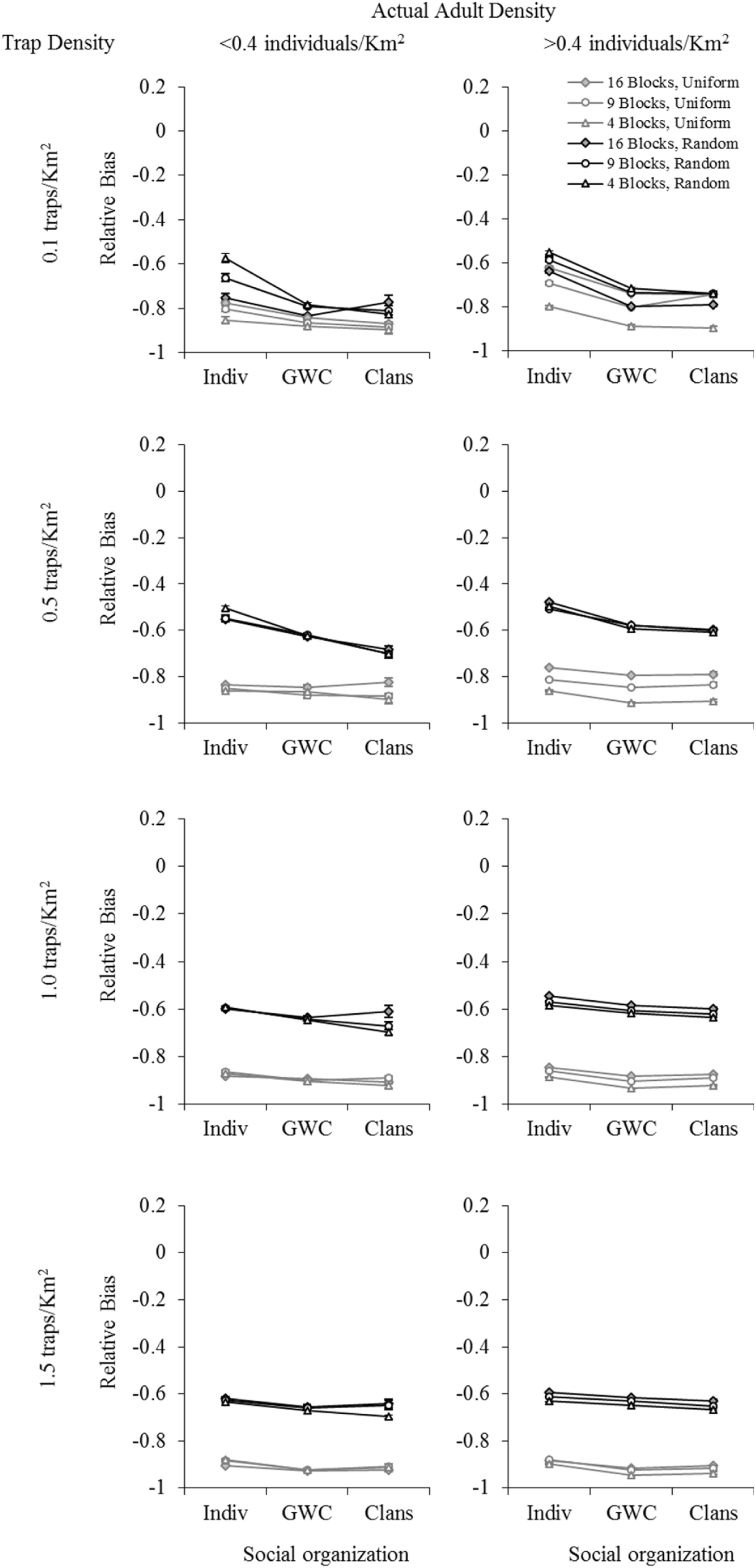
The effect of trap density, sampling scale, social organization, and adult density on mean relative bias in population size estimation using Robust Design with Heterogeneity. Data shown are for a uniform (UTA, grey lines) and random (RTA, black lines) trap arrangement, using actual adult densities. Error bars represent standard errors. B = Blocks (i.e., 16 Blocks, etc.). Social organization: Indiv: non-associating individuals; GWC: groups within clans; Clans: fixed clans.

#### A. POPAN

Trap arrangement explained the majority of variation in relative bias in the five-way fully-factorial ANOVA (Table M in S1 Appendix). We assessed the proportion of variation in the dependent variable explained by a factor or interaction by computing the effect size, *η*^2^ = sums of squares for effect or interaction / total sums of squares. The effect size of the main effect of trap arrangement (*η*^2^ = 0.688) was much higher than those of all other factor main effects and interactions, including interactions involving trap arrangement. All other effects and interactions together explained only a little over 0.2 of the variation in relative bias, while the full ANOVA model accounted for about 0.92 of the total variation in relative bias. Consequently, four-way fully-factorial ANOVAs were performed within each trap arrangement factor level (uniform or random) in order to examine the effects of factors other than trap arrangement.

In general, population size estimates were considerably less biased when a random trap arrangement was used, as compared to when a uniform trap arrangement was used (Fig 2, black vs. grey lines). The greatest bias seen in estimates using a random trap arrangement was about half the magnitude of the smallest bias seen in the case of a uniform trap arrangement (Fig 2; Table A in S1 Appendix). In the case of a random trap arrangement, trap density accounted for much of the variation in relative bias (Table 1; *η*^2^ = 0.518), with higher trap densities tending to yield less biased estimates (Fig 2). The main effect of actual adult density on relative bias was significant (Table 1), but explained only 0.016 of the variation in relative bias (Table 1), suggesting that the adult density experienced in each sampling block did have a small but statistically significant effect on relative bias. There was no significant main effect of sampling scale on relative bias (Table 1), although the largest sampling block size (i.e. 4 blocks) often yielded the most negative values of relative bias (Fig 2). At low trap density (0.1 traps Km^−2^), larger sampling areas (sampling scale of 4 blocks) seemed to give less biased estimates of population density, while at higher trap densities, the smaller sampling scales of 9 or 16 blocks showed slightly less biased estimates (Fig 2). Sampling scale showed significant interactions with trap density (effect sizes ~0.08) and social organization (effect sizes ~0.01). Differences in relative bias among spatial scales were much larger at the lowest trap density than at the three higher trap densities, whereas the fixed clans social organization often yielded the least difference in relative bias among sampling scales (Fig 2). Social organization did have a significant main effect on relative bias (Table 1), but the effect sizes were extremely small (~0.04, Table 1; Figs 4, 5). There were significant two-way interactions between social organization and trap density, sampling scale and adult density (Table 1; Fig 2), but in all these cases the effect sizes were <0.03 (Table 1). Most of the three- and four-way interactions were not significant, and the few that were had effect sizes <0.009.

In the case of uniform trap arrangement, the pattern of effects was somewhat different. Relatively more of the variation in relative bias was explained by sampling scale (Table 1; *η*^2^ = 0.166) and social organization (Table 1; *η*^2^ = 0.291), rather than trap density (Table 1; *η*^2^ < 0.01), even though the main effect of trap density was significant (Table 1). The main effect of actual adult density on relative bias was significant (Table 1), but explained only 0.055 of the variation in relative bias (Table 1). Overall, the smallest absolute values of relative bias were observed when actual adult density in sampling blocks was high, sampling blocks were small in area (i.e., 16 blocks), and there was no social organization into either clans or groups within clans (Fig 2). The two-way interactions between social organization and sampling scale, and between social organization and adult density, were significant, but accounted for only ~0.05 of the variation in relative bias, and three- and four-way interactions were not significant (Tables 1).

#### B. Robust Design

As was the case for the POPAN analyses, for Robust Design, too, trap arrangement explained the majority of variation in relative bias (*η*^2^ = 0.506) in the five-way fully-factorial ANOVA (Table O in S1 Appendix). This was much greater than the variation explained by all other factors and interactions (*η*^2^ < 0.15 for each, 0.46 together). The full ANOVA model accounted for about 0.96 of the total variation in relative bias. As in the case of POPAN, four-way fully-factorial ANOVAs were performed within each trap arrangement factor level (uniform or random), and population size estimates were considerably less biased when a random trap arrangement was used compared to when a uniform trap arrangement was used (Fig 3, black vs. grey lines; S1 Appendix: Table E), although the differences were not as large as those seen in the POPAN analyses. However, compared to the results from the POPAN analyses, absolute values of relative bias were almost always larger in magnitude when Robust Design was used. Even in cases with a random trap arrangement, relative bias values were always more negative than −0.40, as compared to the results from POPAN, where the best combination of factor levels typically gave relative bias values very close to zero.

In the case of a random trap arrangement, there were significant main effects of trap density, spatial scale, social organization, and adult density on relative bias (Table 1). However, the effect sizes of spatial scale and social organization were always <0.06 (Table 1), with a tendency for large sampling block size (4 blocks) and no social organization into clans or groups within clans to give the least relative bias (Fig 3). The largest effect size was for trap density (*η*^2^ = 0.426; Table 1), with the three higher trap densities tending to yield slightly smaller absolute values of relative bias than the lowest trap density of 0.1 traps Km^−2^ (Fig 3; Table E in S1 Appendix). Higher adult female density tended to yield lower absolute values of relative bias (Table 1: *η*^2^ = 0.312; Fig 3). All the two- and three-way interactions among the four factors, although significant, had very small effect sizes (<0.02; Table 1), rendering any further discussion of these interactions uninteresting from a practical point of view. In the case of a uniform trap arrangement, too, there were significant main effects of trap density, spatial scale, social organization and adult density on relative bias (Table 1). However, the only factor with effect sizes >0.1 was actual adult density, with *η*^2^ = 0.498 (Table 1). Again, the trend was for higher adult density to be less biased (Fig 3). Most two-and three-way interactions showed small effect sizes (<0.075).

#### C. Robust Design with Heterogeneity

The overall pattern of results for Robust Design with Heterogeneity was very similar to that for Robust Design, with a slight tendency towards lower relative bias values in the former, in some cases (Fig 4 vs. 3). As was the case for the previous two mark-recapture models, for Robust Design with Heterogeneity, too, trap arrangement explained the majority of variation in relative bias in the five-way fully-factorial ANOVA (Table Q in S1 Appendix) performed using actual (*η*^2^ = 0.682) adult density. This was much greater than the variation explained by all other factors and interactions (*η*^2^ < 0.1 for each). In these ANOVAs, all other effects and interactions together explained about 0.27 of the variation in relative bias, a fraction just slightly greater that that seen in the analyses using POPAN, and considerably less than the fraction of variation explained by all other effects and interactions in the case of Robust Design. The full ANOVA model accounted for about 0.95 of the total variation in relative bias. Four-way fully-factorial ANOVAs performed within each trap arrangement factor level (uniform or random) showed considerably less biased population size estimates when a random trap arrangement was used, as compared to when a uniform trap arrangement was used (Fig 4, black vs. grey lines), although the differences were not as large as those seen in the POPAN analyses. Compared to Robust Design, the differences between relative bias in random versus uniform trap arrangements were of a similar order of magnitude (about 10-31% less bias in the case of random trap arrangement), though slightly larger on an average (Tables E vs. I in S1 Appendix). However, as was the case for Robust Design, compared to the results from the POPAN analyses, relative bias was almost always quite large and negative when Robust Design with Heterogeneity was used. Even in cases with a random trap arrangement, relative bias values were always more negative than −0.47.

In the case of a random trap arrangement, there were significant main effects of trap density, spatial scale, social organization and adult density on relative bias (Table 1). The largest effect sizes were for trap density (*η*^2^ = 0.39: Table 1), with the lowest trap density of 0.1 traps Km^−2^ yielding the highest absolute value of mean relative bias (greatest bias), whereas the second lowest density (0.5 traps Km^−2^) yielded the lowest absolute value of mean relative bias (Fig 4; Table I in S1 Appendix). Social organization also showed a moderate effect size (*η*^2^ = 0.237: Table 1), with the least bias being seen for the case with no social organization into either groups or groups within clans (Fig 4). The effect sizes of spatial scale and adult density were less than 0.064 and two- and three-way interactions also showed small effect sizes (Table 1). In the case of a uniform trap arrangement, too, there were significant main effects of trap density, social organization, and adult density (Table 1). Sampling scale did not show significant main effects (Table 1). However, the only factor with effect sizes >0.1 was trap density (*η*^2^ = 0.347; Table 1), with higher trap density yielding higher absolute values of mean relative bias (Fig 4; Table I in S1 Appendix).

## Discussion

We found that there was a significant main effect of social organization on bias in population estimation in all our analyses, but the effect sizes were usually small. We found that arranging traps randomly in the sampling blocks tended to yield relatively less biased estimates of population size, regardless of the mark-recapture model used (Figs 2–4). All three mark-recapture methods typically underestimated population size, often by a large degree (Figs 2–4), and POPAN clearly gave less biased estimates than the two Robust Design models we tested (Fig 2 vs. Figs 3, 4). Moreover, the combinations of random trap arrangement with relatively high trap and adult densities, for which POPAN gave more or less unbiased estimates, were also the combinations wherein social organization did not markedly affect bias in population size estimation (Fig 2). In this section, we discuss these major findings in the context of study design for population size estimation in wild animals more generally, and in Asian elephants, specifically. Finally, we conclude by discussing some of the broader studies possible using our simulation framework.

The most reassuring result from our study was that, although there were effects of social organization on bias in population estimation, it could be mitigated using random trap arrangement, high trap density, and appropriate spatial scale of sampling. When we used POPAN, which performed better overall compared to the other methods, social organization and spatial scale of sampling explained a moderate amount of variation in relative bias in the case of a uniform trap arrangement (Fig 2). Under high adult density, uniform trap arrangement, and analysis based on POPAN, the relative bias of the fixed clans social organization almost doubled compared to that of non-associating individuals, while random trap arrangement resulted in almost no difference in relative bias based on social organization (Fig 2). A potential reason for the fixed clans and groups within clans cases producing more biased results compared to the non-associating individuals case under a uniform trap arrangement could be that, in the first two cases, the number of individuals with very low or zero capture probabilities increased, thus increasing the variance in capture probabilities which would result in biased estimates of population size. Such an increase could be a consequence of more individuals falling in areas not covered by the traps because of the aggregation of individuals in the fixed clans and groups within clans social organization. The problem may be exacerbated in the case of uniform traps compared to random traps if groups end up at the edges of sampling blocks because the uniform trap arrangement does not cover sampling block edges. The social organization of individuals, into clans or groups within clans as compared to non-associating individuals, did not markedly affect the value of relative bias under the combination of factor levels (random trap arrangement and relatively high trap and adult densities) for which POPAN yielded more or less unbiased estimates of population size (Fig. 2). This result clearly suggests that if one can ensure a reasonably high density random trap arrangement in a study aimed at estimating population size in a social species, then one can be reasonably confident of obtaining unbiased estimates using POPAN.

The random trap arrangement always yielded more unbiased estimates than the uniform trap arrangement in our study, except occasionally under low trap density (see Figs. 2–4). This was probably because random trap arrangement ensures a better overall coverage of a localized area than uniform trap arrangement, especially when movement of individuals is random, as it was in our simulations. In real situations where movement is highly nonrandom, for example along trails or in the vicinity of resource patches, it may help to concentrate the sampling effort in areas of high movement [44–46], although a random arrangement of traps within a sampling block in those areas may still be better than a uniform trap arrangement. While we have not studied this so far, our simulation framework can be used to examine the reliability of convenience sampling in the future. Sampling scales were of the order of the size of clan home ranges in our simulations. It is possible that a random trap arrangement might not perform as well for species with smaller home ranges because, in such cases, entire home ranges may fall into ‘holes’ (see below) where the sampling effort is zero.

It is not surprising that relatively high trap density and adult density yielded less biased estimates of population size in our study. It is known that low trap densities can result in the formation of areas with no trapping effort called ‘holes’, resulting in some individuals having reduced or even zero capture probabilities, thus inducing bias in superpopulation size estimates [8,45]. Ultimately, trap density and population density in the sampling blocks need to be commensurate to ensure relatively unbiased estimates of population size. If population density is low, capture probabilities need to be relatively high in order to obtain reliable estimates of population size [11,47]. This, unfortunately, creates a catch-22 situation because it implies that one must have at least an estimate of local population density to decide upon an efficacious trap density for a study. However, crude rules of thumb based on rough estimates of overall population density can be used in the absence of detailed prior knowledge of local population densities. In an individual-based simulation study based on data from grizzly bears (*Ursus arctos horribilis*), Boulanger *et al.* [11] found that they obtained reliable estimates only for populations with a real size above 50, and that capture probabilities above 0.2 greatly improved the estimates. Similarly, Otis *et al.* [47] suggested that populations of size around 50 may require capture probabilities as high as 0.4-0.5 to obtain reliable mark-recapture estimates, whereas even capture probabilities of ~0.2 yield reliable estimates when the real population size is 200. In fact, individual-based simulations suggested that both uniform and random trap arrangements can yield fairly unbiased mark-recapture estimates of population size, as long as movement per time-step is small, trap density is really high and sampling is done over at least 20 occasions [46].

We next discuss possible reasons for why almost all combinations of factor levels yielded underestimates of population size, regardless of mark-recapture model, and why POPAN outperformed the two Robust Design models in our study. The consistent underestimation of population size is most likely explained by the heterogeneity in capture probability among individuals. Individuals with higher capture probabilities will be more likely to initially get ‘marked’ and subsequently recaptured, such that later capture cohorts will show a proportion of marked individuals that is greater than the actual proportion in the population, thereby resulting in underestimation of population size [48]. The high absolute values of relative bias in population size estimates using both Robust Design and Robust Design with heterogeneity indicate a basic limitation of these models when used on the kind of datasets that our simulations generated. The most likely explanation for the poor performance of Robust Design methods is that they estimate population size within each primary interval (seven days in our study), whereas POPAN uses all the data across the primary intervals (13 weeks in our study). Consequently, there was a big difference in sampling effort used for any particular estimate in our study, with the sampling effort for POPAN being substantially higher than for either of the two Robust Design models. Moreover, the Robust Design estimator is quite sensitive to permanent emigration from the study area during the primary intervals [8,47,49], an occurrence that was possible in our simulations. The above considerations suggest that if Robust Design methods are to be used for studies structured in a manner similar to our simulations, then either longer primary intervals should be used, or the intensity of sampling in the secondary intervals should be much increased, in order to obtain relatively unbiased estimates of population size. Unfortunately, longer primary intervals would automatically make it more likely that the assumption that the population is closed during primary intervals will be violated. Another result in our study was that relative bias did not always decrease with increase in trap density in the Robust Design analyses. This could happen if an increase in trap density disproportionately increased the capture probability of only some individuals in the study population. Such a situation could arise if some individuals permanently emigrated from the population (sampling block in our case), thus making their recaptures impossible within any particular primary interval. It is interesting to note that the inclusion of variation in capture probabilities in the Robust Design with heterogeneity model did not substantially reduce the bias in population size estimates as compared to the Robust Design model without heterogeneity, although such a result has been observed previously also [11]. This indicates that the real problem with the Robust Design models in our study was probably the low sampling effort per primary interval. In actual studies on real populations, this limitation will be hard to circumvent due to logistical constraints, suggesting that POPAN may generally be a better option for mark-recapture estimation of population size for wide-ranging species.

Overall, our study reinforces several recommendations about the design of mark-recapture studies that have been already made before. For example, that (a) closure of the study area be ensured when using closed-capture estimators [8,47,49], (b) trap densities be high enough so as to minimize holes [8,45], (c) more intense sampling be done in the case of small populations, to compensate for intrinsically low capture probabilities [11,47], and (d) individual heterogeneity be adjusted for, to a degree, by using estimators that explicitly account for heterogeneity [43]. Moreover, we also show that non-independence in movement between individuals can increase the variance in individual heterogeneity and, thus, additional efforts in sampling and trap array design are needed to ensure unbiased estimates in the case of social species. Our results also strongly suggest that uniform trap arrangements should be avoided when the available trap densities do not provide adequate coverage and when the study species is social. This is similar to an earlier suggestion that, if trap density is low, then the study area should be divided into smaller units and traps should be set in a randomly chosen set of sub-areas in each sampling occasion to ensure better coverage of localized areas [8]. In the case of social species with very large home ranges, like the Asian elephant, for which large study areas are required, a hybrid of these two techniques, i.e. random placement of traps in randomly chosen sub-areas across sampling periods, may possibly give the best results. Trap densities used in previous studies of relatively large ranging species, for example photographic capture-recapture studies on the Bengal tiger *(Panthera tigris tigris*) [50], have been in the range of 0.2-0.7 traps/Km^2^. This is close to the second trap density factor level (0.5 traps/Km^2^) in our study and we obtained reasonably unbiased estimates (averaged bias ~ 5% in case of POPAN) at this trap density when using a random arrangement of traps. In the case of species with very large home ranges, like the Asian elephant, our results indicate that logistical constraints are likely to preclude the use of the Robust Design estimator in a large study area, as it would be very difficult to obtain enough resources to ensure a large sampling area along with high sampling intensity during long enough sampling periods, while keeping the population geographically closed. For such species, POPAN appears to be a much better option for mark-recapture studies. We tried to mimic a long-term study situation in our simulated trapping protocol and, thus, we sampled on 84 occasions in each sampling block in each replicate simulation. Real studies usually sample for much shorter periods, and a good method to determine an adequate number of sampling occasions would be to look for plateauing of the cumulative uniquely identified/marked individuals curve over the study period [35]. Our results also reinforce the view that reliable estimates of population size for a wide ranging social species like the Asian elephant can only be obtained by carefully planning the study design. Surveyors need to ensure sampling of an area which is several times the size of the average home range of the species [51], with traps either randomly placed or placed to minimize the perimeter to area ratio, in randomly chosen sub-areas of the study area [8] if sampling the entire area at one go is not feasible due to practical constraints. To ensure high capture probabilities, trap densities should definitely be higher than 0.1 traps/Km^2^, preferably closer to 0.5 traps/Km^2^, or enough sampling occasions should be included in the study to ensure thorough sampling of the study population. Spatial capture-recapture models [52] may provide an alternative, using which relatively unbiased population size estimates may be obtained while sampling smaller areas [53]. Spatial capture-recapture models have been found to perform well under a variety of trap arrangements and animal movement patterns [53]. They might be useful in estimating elephant abundance too, and it might be worth examining the effect of social organization on such models in the future.

In conclusion, we would like to stress that, although our present study was parameterized with specific reference to southern Indian populations of Asian elephants, the simulation framework we have developed has far broader applications. The simulation framework can be altered by using species-specific parameters to assist with the design of future mark-recapture studies of Asian elephants and other wide ranging and/or social species. Some marine mammals move in social groups and occupy large home ranges, and may provide similar challenges as those faced in estimating elephant abundance. Uniformly spaced sampling, which provided biased estimates in our study, has been used in the estimation of abundance of several whale species and the bottlenose dolphin [54–56]. Robust Design analysis has been used to estimate the abundance of dolphin species [57], although capture probabilities in that study were larger than 0.1 (which was the capture probability in our study). Mark-resight models that accommodate detection heterogeneity have been used to estimate population size of ungulates [58,59], although these models assume population closure and may not be applicable to more wide-ranging species. Our simulation framework can be modified and used to assess relative bias under the circumstances specific to studies such as those mentioned above, and to find out whether different sampling designs would help improve these abundance estimates. Observational studies on the behaviour and ecology of social animals in the wild are typically beset with the problem of an inherent lack of replication, making it difficult to make even somewhat tentative inferences about causal ecological factors affecting aspects of behaviour. The simulation framework described in this paper can be modified to examine problems that go well beyond those of reliably estimating population size, such as how spatial distribution of resources and movement constraints due to topography influence social organization, how adopting different criteria for defining an association between individuals can affect the social organization that will be inferred from observed associations, or what kind of sampling strategy and effort will be optimal for assessing various aspects of social organization and behaviour. In our present study, we simulated movement of individuals that was random, except to the extent that there was a tendency for coordinated movement among individuals belonging to various levels of social organization. This simulation framework can be extended to accommodate non-random movement due to different spatial distributions of resources and/or topographical features. Similarly, it is also possible to expand this simulation framework to incorporate specific kinds of behavioural interactions among individuals belonging to different hierarchical levels of social organization. Such simulations will provide us with a platform to address various questions regarding the effects of movement, density and resource distribution on social organization, as well as the consequences of such factors for the optimal design of sampling strategies in observational studies of social behaviour and ecology in wild populations.

## Methods

### Simulation

The simulation was programmed and run using MATLAB^®^ R2011a [60]. Only adult females were considered in the simulation, as only female Asian elephants, and not males, exhibit coordinated movement among themselves [20,21]. A study in Nagarahole National Park had identified 37% of all individuals to be adult females [61]. For the purpose of this paper, we consider individuals >=15 years of age as adults, as treated in Arivazhagan and Sukumar [61]. Subsequent long-term monitoring of individually identified and aged elephants in the Nagarahole and Bandipur National Parks [62] showed the percentage of adult females to be about 35% (Nandini Shetty, Keerthipriya P, Vidya TNC, unpublished data). The total density of elephants in the Nilgiris-Eastern Ghats Reserve had been estimated at 0.5-0.83 elephants Km^−2^ [63], and that within Nagarahole National Park at 2.25 [63] and 3.3 elephants Km^−2^ [27]. Thus, the density of adult females across the region could range from about 0.18-0.3 (Nilgiris-Eastern Ghats Reserve) to 0.79-1.16 Km^−2^ (Nagarahole National Park). Accordingly, we chose four different adult female density regimes of 0.2, 0.4, 0.8, and 1.2 adult females Km^−2^ for the simulations. It has been shown earlier that Asian elephant clans in southern India have home ranges of about 530-670 Km^2^ [21,64]. Consequently, we obtained clan home range sizes for the simulations from a close-to-continuous uniform distribution between 500-700 Km^2^, and assigned them randomly to clans. The total study area in the simulations was set to be 2500 Km^2^ (50 Km × 50 Km). This area was further divided into 4, 9, or 16 square sampling blocks and sampling was done independently within each sampling block. Trap locations within each sampling block were either arranged in a uniform square grid, or chosen randomly, following a close-to-continuous uniform random distribution. In both cases, trap densities of 0.1, 0.5, 1.0, and 1.5 traps Km^−2^ were used in different simulations. Three different social organizations were investigated: fixed clans, groups within clans, and non-associating individuals. The fixed clans case had only one level of social organization, wherein individuals associated with one another primarily within clans, regardless of group. The groups within clans case had two levels of social organization, in which individuals within a group primarily associated with one another, but could also associate with lower probabilities with individuals from other groups within the same clan. In the non-associating individuals case, individuals moved in a more or less randomly chosen direction, without any greater association between individuals from the same group or clan. The various cases (factor levels) for each of these factors are listed in Table 2. Combinations of factor levels were used to run simulations in a fully factorial, randomized cross design. Ten replicates were run for each simulation with a unique combination of factor levels, resulting in a total of 2880 simulations (2 trap arrangements × 4 trap densities × 3 sampling scales × 3 social organizations × 4 adult female density regimes × 10 replicates).

**Table 2.** Different cases (levels) of parameter values used in the simulations for the factors trap arrangement, trap density, sampling scale, social organization and overall adult female density. The various simulations used all combinations of factor levels, crossed amongst themselves.

| **Parameter** |  | **Case 1** | **Case 2** | **Case 3** | **Case 4** |
| --- | --- | --- | --- | --- | --- |
| **Trap Arrangement** |  | Uniform square grid | Uniformly random | - | - |
| **Trap Density** |  | 0.1 traps/Km<sup>2</sup> | 0.5 traps/Km<sup>2</sup> | 1 trap/Km<sup>2</sup> | 1.5 traps/Km<sup>2</sup> |
| **Sampling Scale** |  | 16 blocks | 9 blocks | 4 blocks | - |
| **Social Org. (AP range)** | **Group-mates** | 0.5-0.7 | 0.5-0.7 | 0-0.001 | - |
|  | **Clan-mates** | 0-0.2 | 0.5-0.7 | 0-0.001 | - |
|  | **Other inds.** | 0-0.001 | 0-0.001 | 0-0.001 | - |
| **Adult Density** |  | 0.2 individual/Km<sup>2</sup> | 0.4 individuals/Km<sup>2</sup> | 0.8 individuals/Km<sup>2</sup> | 1.2 individuals/Km<sup>2</sup> |

For social organization, Case 1 is groups within clans, Case 2 is fixed clans and Case 3 is non-associating individuals. AP = association probability.

A multi-level social organization thought to be reasonably reflective of Asian elephant populations when we started this work [20,25,65,] was implemented. The total number of groups in a population was set to be one-fifth of the total number of individuals, and the total number of clans to be one-sixth of the total number of groups (based on the limited information available about clan size; see 20,65). Each group was assigned a group size drawn randomly from a normal distribution with mean 5.0 and standard deviation of 2.33. The group size was then rounded off to the nearest integer and these assigned group sizes were then summed up over all groups to yield the total population size of adult females. The stochasticity inherent in this procedure for assigning group sizes sometimes resulted finally in a population size slightly different from that initially assigned based on the specific adult female density used in that particular simulation and, consequently, the new population size obtained by summing up all the group sizes was used for all further analyses. The number of groups in each clan was picked from a uniform integer distribution ranging from 4 to 8, and this step was repeated until the total number of groups equalled the original number. Subsequently, each individual was accordingly assigned a group and a clan identity. The home range centres for each clan were assigned at random within the 2500 Km^2^ study area. The home range area for each clan was then assigned by randomly picking areas from a close-to-continuous uniform random distribution between 500-700 Km^2,^ and a length-to-breadth ratio from a close-to-continuous uniform random distribution between 0.5 and 2. One random location was then chosen within each clan's home range (and designated the clan centre), and each group within that clan was assigned an initial position located randomly within 2.5 Km of the clan centre. All individuals in each group were then randomly assigned initial positions within 500 m of their respective initial group positions.

The different social organizations were implemented through different patterns of pair-wise association probabilities (APs) between individuals from the same group, from different groups within the same clan, or from different clans, respectively. APs quantified the probability that two individuals moved towards each other when in close proximity (within 2 km, see [66]). AP values used for individuals of the same or different groups and clans in order to define the three social organizations studied are listed in Table 2. Pairs of individuals were categorised based on their respective group and clan memberships, i.e., individuals belonging to the same group, individuals belonging to the same clan but different groups, and individuals belonging to different clans, were given different, or similar, pair-wise AP values, based on type of social organization (Table 2). The patterns of AP values were set up such that, in the groups within clans case, individuals would most often associate with group-mates, second most often with non-group clan-mates, and least often with individuals from other clans. For the fixed clans case, it was ensured that individuals would associate equally often with all clan-mates, irrespective of group, and would associate much less frequently with individuals from other clans, thereby effectively eliminating group as an hierarchical level in the social organization. For the non-associating individuals case, the APs were identical, and extremely low, between all pairs of individuals, regardless of group or clan identity. In every simulation, each pair of individuals was assigned an AP value chosen randomly from the applicable range of values as listed in Table 2. This was done by randomly picking an AP value from a close-to-continuous uniform distribution spanning the relevant AP range, depending on the type of social organization being used for that simulation and the group and clan identities of those two individuals.

Each simulation consisted of individuals being moved over 600 simulated days in time steps of 2 hours, i.e., 12 × 600 time steps. Within each time step, each individual was taken as the focal individual sequentially, and all individuals within its sensory range, i.e., 2 Km [60], were identified. Among these individuals within 2 Km of the focal individual, one individual was stochastically chosen according to a probability proportional to its AP with respect to the focal individual, after the sum of the APs of all those individuals with respect to the focal individual had been normalized to one. The focal individual was then either moved towards the chosen individual with a probability equal to their AP, or it was moved in a randomly chosen direction with probability 1-their AP. The movement of the focal individual outside its home range was prevented by moving it towards the centre of the home rage if it did exit the home range when moved. All individuals were assumed to move with a constant speed of 5 Km/day (~0.42 Km/time step). Each time step in a simulation was concluded when the above procedure had been successfully run for all individuals; all individuals were moved to their new positions simultaneously at the end of the time step (See Figure D in S1 Appendix for snapshots of the simulation).

In field studies attempting population size estimation, typically only a part of the range of the populations being studied is actually sampled [27,30,35]. Consequently, we also wanted to assess the effect of the relative proportion of the total study area sampled on bias of mark-recapture estimates of population size. Therefore, in different simulations, the total study area was divided up into either 16, 9 or 4 square sampling blocks of equal area, thus representing sampling efforts corresponding to 1/16, 1/9 or 1/4 of the total study area, respectively. All such blocks were sampled in each replicate simulation. Sampling traps in any given simulation were either arranged in a uniform square grid in all the blocks, or were arranged randomly in all the blocks by drawing trap locations from a uniform distribution. In each simulation, sampling was done for every 3^rd^ week of every 30-day month (15^th^ day to 21^st^ day of each month), from the 251^st^ day onward to the 600^th^ day. In initial trial simulations, we often found that there were changes in the level of spatial dispersion shown by groups within clans during the initial stages of the simulation, but that the dispersion patterns appeared to stabilize with time. Consequently, in the simulations for the main study, the first 250 days were left out in order for the mean clan hull-area to reach a plateau, thereby avoiding the possibility of transient behaviour artifactually affecting the results of how the various factors in our study affected bias of mark-recapture estimates of population size. On each sampling day, sampling was done for the first six time steps (equalling half a day). In each of these time steps, all individuals within 100 m of the various trap locations in a given sampling block were recorded as captures within that sampling block. For each sampling block, data on captures were stored as capture histories: matrices in which each row represented an individual and each column represented a sampling day. Any individual captured one or more times during the 6 time steps in a day, at any of the trap locations within a given sampling block, was recorded in the capture history as a single capture for that day in that sampling block. This was done so that the number of sampling occasions usually remained less than the total number of individuals ever captured as this is needed to get a good fit while fitting the models. Thus, each matrix had 0's and 1's as entries, representing the absence or capture of an individual, respectively, in that sampling block for that day. These capture histories of individuals that were “trapped” in each sampling block were used for mark-recapture analyses. In order to find the actual number of individuals present, we also recorded from the simulations the total number of unique individuals located in that sampling block in any of the six time steps constituting that sampling day, regardless of their distance from trap locations. In each simulation, the total number of unique individuals located in a sampling block was recorded and stored for (a) each sampling week, (b) the entire sampling duration, i.e., 12 sampling weeks, and (c) the entire duration of the simulation, i.e., 350 days (after leaving out the first 250 of 600 days). These values were used as real population size values in the analysis of bias in population size estimates obtained from mark-recapture analyses using either the POPAN or Robust Design (with and without heterogeneity) models. The authors may be contacted for the codes used to implement the simulations. Zero estimates were removed from the data as they would give an error when used in the relative bias formula. Estimates with abnormally high relative bias values (>5.0) were also removed as they were usually obtained due to bad model fitting. Since we monitored only adult females, we did not take demographic variables into account in our simulations. The number of adult females is not expected to change within the period of a year as mortality in adult female Asian elephants is very low [67]. Females may calve during the year, but calves are not counted in this study. Subadult females surviving to enter the adult female category during the subsequent year would also not be considered given our study duration.

### Analyses

All the mark-recapture analyses were done using the POPAN and Robust Design models implemented using RMark [68], which is an R library providing an interface between Program MARK [69] and the R programming language [70]. Altogether, three models (POPAN, Robust Design without heterogeneity and Robust Design with Heterogeneity: two mixtures) were tested for bias as a result of various combinations of trap arrangement, trap density, adult density, sampling scale, and social organization.

The POPAN estimator [38] models the capture of individuals using three parameters: survival probability (*φ*), capture probability (*p*), and the probability of entry into the study area from the superpopulation (*pent* or *b*). The superpopulation (*N*) in this model comprises of all individuals that ever enter the study area during the sampling period and is obtained using maximum-likelihood estimation [69]. In our analyses, we set the probability of survival to be constant over time as there were no demographic processes, i.e., births and deaths, included in the simulation. However, the survival probability was not set to one because permanent emigration is treated as a death in this model. The probabilities of capture and entry were also set to be constant in our simulations because the movement of groups (or individuals when they were not associating with each other) was uniformly random over time. The population size was estimated using all sampling occasions and the three-week gaps in sampling were explicitly defined in the model [69].

Robust Design methods [39–41] incorporate the notions of both open and closed populations while modelling mark-recapture, and captures are modelled in a hierarchical fashion. In this family of methods, there are several different sets of captures employed, during each of which the population is assumed to be closed, whereas the population is considered to be open in the time between the capture sets. These sets are called primary occasions, while the capture occasions within them are called secondary occasions. The Robust Design with Heterogeneity model [42,43] tries to account for variation in capture probabilities among individuals by modelling the population as consisting of several groups of individuals (mixtures), with each group having a different capture probability that is, nevertheless, the same among all members of a group. It has been shown that a Closed-Captures model with heterogeneity that assumes two mixtures is parsimonious and yields relatively unbiased and precise estimates [43]. We used Robust Design without heterogeneity as well as Robust Design with heterogeneity, with two mixtures, in our study. The parameters used in Robust Design models are survival probabilities (*φ*), capture probabilities (*p*), probability of immigration (1-*γ*′), probability of emigration (*γ*″) and population sizes for each primary occasion (*N*). A Robust Design with Heterogeneity model, using two mixtures with different capture probabilities, has another parameter (*π*) which is the proportion of individuals in the first mixture. In our analyses, survival probability was set to be constant over time (as in POPAN analysis) as demographic processes were not included in our simulations. The capture probability, and the probabilities of emigration and immigration were set to be constant over time as the movement of groups, or individuals when they were not associating with each other, was uniformly random in our simulations. Constant probabilities of emigration and immigration model the movement into and out of the study area as being random [40,41]. The proportion of individuals in the first mixture (*π*) was also set to be constant. There were 12 primary occasions (the 3^rd^ week of each month), consisting of seven secondary occasions each (the seven days in the 3^rd^ week of each month), in our analyses. The Robust Design models we used then estimated population size in each sampling block for each primary occasion of sampling.

As mentioned above (Simulation), in our simulations, we modelled three different social organizations by assigning different patterns of APs, reflecting the pair-wise likelihood of coordinated movement between individuals, to individuals from the same group, from different groups within the same clan, and from different clans. In order to assess whether these three patterns of APs actually resulted in the three desired social organizations (groups within clans, only clans, non-associating individuals), we calculated pair-wise association indices (AIs) for all pairs of individuals within all three social organizations for the first 350 days in multiple simulations involving the different social organizations, although we present data from only a representative set of simulations (Fig 1). Pair-wise AIs [71] are extensively used in studies of social organization in animals and are calculated as the proportion of sightings in which two individuals are seen associating with each other with respect to the total number of sightings in which the two individuals are seen. In our simulations, we performed *K*-means clustering [72,73] within each clan, based on spatial locations of individuals, and the optimum *K* was found using cluster silhouettes [74]. Two individuals were deemed to be associating if they were found to be in the same cluster. The associations determined by the clustering were then used to compute pair-wise AIs, and the distribution of pair-wise AIs was plotted within clans, and also within groups, for seven consecutive 50 day periods for the first 350 days of the simulations, to ascertain whether our simulations of movement with three different patterns of APs actually resulted in three detectable social organizations of the desired kind.

For all estimates of population size in each sampling block in each simulation, relative bias was calculated as

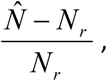

where *N_r_* is the real population size of the population sampled and *N̂* is the estimated population size for the same population. For the final analysis of how relative bias of population size estimates obtained from each of the three models (POPAN, Robust Design with and without heterogeneity) was affected by the factors trap arrangement, trap density, adult density, sampling scale, and social organization, we carried out five-way fully factorial analyses of variance (ANOVAs), taking the mean relative bias across all sampling blocks for each replicate simulation as the dependent variable, and treating the five factors as fixed and crossed among themselves. In these analyses, the factor adult density was taken as reflecting the four different levels of initial adult density (0.2, 0.4, 0.8, and 1.2 individuals Km^−2^), one of which was set as the initial value across the entire study area at the beginning of each simulation. Thus, simulations were classified with regard to adult density based on which initial density was used in that simulation, and these ANOVAs use the factor initial density, with four levels. The reason for wanting to treat adult density as a factor in the analyses was that we wanted to ascertain whether degree of bias in mark-recapture estimates of population size was affected by the overall population density in the study area. In all our simulations, the recording of real population size and the calculation of estimated population size were done for individual sampling blocks. Upon examining the results of the simulations, it became apparent that often the average (over time) adult density in a sampling block exhibited considerable variation among sampling blocks within the same simulation with a specific initial adult density. Consequently, we decided to do a separate set of five-way ANOVAs that would reflect the effects of the actual mean adult density, observed over time in a sampling block, on bias in population size estimation. In order to do this, we recorded the average number of individuals present per day, over the entire simulation (350 days), for each sampling block in each simulation. Dividing this average by the area of the sampling block yielded the mean actual adult density in each sampling block. Next, we examined all the sampling blocks for the first replicate of each simulation, regardless of which level of initial adult density was used in the simulation. Based on the median adult density observed across sampling blocks over all the simulations, we classified all sampling blocks from the first replicate of each simulation into two categories: those with adult density >0.4 Km^−2^ and those with adult density <0.4 Km^−2^. The same process was repeated for all ten replicates of each type of simulation. Using the median adult density across sampling blocks as the cut-off for this categorization ensured roughly similar numbers of sampling blocks in each category. Thereafter, the relative bias was averaged across all sampling blocks in each of these two categories, and that was used as the dependent variable in the ANOVA, with actual adult density as a factor with two levels (<0.4 Km^−2^ and >0.4 Km^−2^). The other four factors in the ANOVAs remained the same as described previously, i.e., trap arrangement, trap density, sampling scale and social organization. Thus, at the end of this procedure, for each combination of levels of the other four factors, there were ten replicate values each of mean relative bias for the two levels of the factor actual adult density (data shown in S2 Appendix).

Because we treated adult density in two different ways, a total of six five-way ANOVAs were carried out. In all these ANOVAs, we found that trap arrangement accounted for a large proportion of variation in relative bias, compared to all other factors and interactions (Tables M, N in S1 Appendix). Consequently, in order to better tease apart the effects of other factors, we also performed four-way fully-factorial ANOVAs within each level of trap arrangement (uniform or random) for each model, and for both initial and actual adult female density, resulting in a total of twelve four-way ANOVAs. All ANOVAs were performed using Statistica version 5.0 [75]. All pair-wise multiple comparisons were done using Tukey’s HSD test [75]. The real superpopulation size for the POPAN analysis was taken as the total number of unique individuals present in the sampling block for the entire sampling duration. The real population size for Robust Design was taken as the total number of unique individuals present in a block for a particular primary sampling occasion. Results from only the analysis using actual adult density as factor are reported in the Results section. Please see S1 Appendix for results from the analysis using initial adult density as a factor.

## Author contributions

TNCV conceived of the study, TNCV, MG and AJ designed the simulations and analyses, MG and TNCV implemented the simulations, MG did the analyses, TNCV, MG and AJ wrote the manuscript.

## Acknowledgments

We thank T.N.C. Anand, Indian Institute of Technology Madras, for very helpful discussions regarding the early formulations of the simulation model and for help with the basic programming framework, upon which the simulation model was developed.

## S1 Appendix

Results of analyses of the effects of trap arrangement, trap density, sampling scale, social organization and adult density on relative bias in population size estimation using three mark-recapture estimators. This appendix contains ANOVA tables for all four- and five-way ANOVAs, as also tables listing the mean relative bias for various combinations of factor-levels, along with the pattern of significance among them in pair-wise multiple comparisons. Mean values and significance levels in pairwise comparisons are shown only for main effects and interactions that had a significant effect in the ANOVA.

## S2 Appendix

Data for all the statistical analyses carried out.

## References

1. Kingsland SE. Modeling nature: episodes in the history of population ecology. 2^nd^ ed. Chicago: University of Chicago Press; 1995.

2. Turchin P. Complex population dynamics: a theoretical/empirical synthesis. Princeton: Princeton University Press; 2003.

3. Clutton-Brock TH. Sex ratio variation in birds. Ibis. 1986; 128: 317–329.

4. Clutton-Brock TH, Iason GR. Sex ratio variation in mammals. Q Rev Biol. 1986; 61: 339–374.

5. Jones JP, Asner GP, Butchart SH, Karanth KU. The ‘why’, ‘what’ and ‘how’ of monitoring for conservation. In: Macdonald DW, Willis KJ, editors. Key topics in conservation biology 2. Oxford: John Wiley & Sons; 2013. pp. 327–343.

6. Nunney L, Elam DR. Estimating the effective population size of conserved populations. Conserv Biol. 1994; 8: 175–184.

7. Frankham R. Effective population size/adult population size ratios in wildlife: a review. Genet Res. 1995; 66: 95–107.

8. Williams BK, Nichols JD, Conroy MJ. Analysis and management of animal populations. San Diego: Academic Press; 2002.

9. Amstrup SC, McDonald TL, Manly BF. Handbook of capture-recapture analysis. 1st ed. Princeton: Princeton University Press; 2005.

10. Seber GAF. The estimation of animal abundance and related parameters. New York: MacMillan Press; 1982.

11. Boulanger J, McLellan BN, Woods JG, Proctor MF, Strobeck C. Sampling design and bias in DNA-based capture-mark-recapture population and density estimates of grizzly bears. J Wildl Manage. 2004; 68: 457–469.

12. McCullagh P, Nelder JA. Generalized linear models. New York: Chapman & Hall/CRC Press; 1989.

13. Schmutz JA, Ward DH, Sedinger JS, Rexstad EA. Survival estimation and the effects of dependency among animals. J Appl Stat. 1995; 22: 673–682.

14. Cox DR, Snell EJ. Analysis of binary data. New York: Chapman & Hall/CRC Press; 1989.

15. Bishop CJ, White GC, Lukacs PM. Evaluating dependence among mule deer siblings in fetal and neonatal survival analyses. J Wildl Manage. 2008; 72: 1085–1093.

16. Kummer H. Primate societies: group techniques of ecological adaptation. Chicago: Aldine; 1971.

17. Dunbar R. Primate social systems. New York: Cornell University Press; 1988.

18. Aureli F, Schaffner CM, Boesch C, Bearder SK, Call J, Chapman CA, et al. Fission-fusion dynamics: new research frameworks. Curr Anthropol. 2008; 48: 627–654.

19. McKay GM. Behavior and ecology of the Asiatic elephant in southeastern Ceylon. Smithson Contrib Zool. 1973; 125: 1–113.

20. Sukumar R. The Asian elephant: ecology and management. Cambridge: Cambridge University Press; 1989.

21. Baskaran N, Balasubramanian M, Swaminathan S, Desai AA. Home range of elephants in the Nilgiri Biosphere Reserve, south India. In: Daniel JC, Datye HS, editors. A week with elephants: proceedings of the international seminar on the conservation of Asian elephant. Bombay: Bombay Natural History Society and Oxford University Press; 1995. pp. 296–313.

22. Fernando P, Lande R. Molecular genetic and behavioral analysis of social organization in the Asian elephant (*Elephas maximus*). Behav Ecol Sociobiol. 2000; 48: 84–91.

23. Vidya TNC, Sukumar R. Social organization of the Asian elephant (*Elephas maximus*) in southern India inferred from microsatellite DNA. J Ethol. 2005; 23: 205–210.

24. Shetty N, Keerthipriya P, Vidya TNC. Group size constraints may mask underlying similarities in social structure: a comparison of female elephant societies. BioRxiv doi: https://doi.org/10.1101/099614.

25. de Silva S, Ranjeewa AD, Kryazhimskiy S. The dynamics of social networks among female Asian elephants. BMC Ecol. 2011; 11: 17.

26. Dawson S. A model to estimate density of Asian elephants (*Elephas maximus*) in forest habitats. M.Sc. Thesis, University of Oxford. 1990.

27. Karanth KU, Sunquist ME. Population structure, density and biomass of large-herbivores in the tropical forests of Nagarahole, India. J Trop Ecol. 1992; 8: 21–35.

28. Varman KS, Ramakrishnan U, Sukumar R. Direct and indirect methods of counting elephants: a comparison of results from Mudumalai Sanctuary. In: Daniel JC, Datye HS, editors. A week with elephants: proceedings of the international seminar on the conservation of Asian elephant. Bombay: Bombay Natural History Society and Oxford University Press; 1995. pp. 331–339.

29. Baskaran N, Sukumar R. Karnataka elephant census 2010. Technical report to the Karnataka Forest Department. Bangalore: Asian Nature Conservation Foundation and Indian Institute of Science; 2011.

30. Goswami VR, Madhusudan MD, Karanth KU. Application of photographic capture–recapture modelling to estimate demographic parameters for male Asian elephants. Anim Conserv. 2007; 10: 391–399.

31. Williams AC, Johnsingh AJT, Krausman PR. Population estimation and demography of the Rajaji National Park elephants, North-West India. J Bomb Nat Hist Soc. 2007; 104: 145–152.

32. de Silva S, Ranjeewa AD, Weerakoon D. Demography of Asian elephants *(Elephas maximus)* at Uda Walawe National Park, Sri Lanka based on identified individuals. Biol Conserv. 2011; 44: 1742–1752.

33. Hedges S, Johnson A, Ahlering M, Tyson M, Eggert LS. Accuracy, precision, and cost-effectiveness of conventional dung density and fecal DNA based survey methods to estimate Asian elephant (*Elephas maximus*) population size and structure. Biol Conserv. 2013; 159:101–108.

34. Gray TN, Vidya TNC, Potdar S, Bharti DK, Sovanna P. Population size estimation of an Asian elephant population in eastern Cambodia through non-invasive mark-recapture sampling. Conserv Genet. 2014; 15: 803–810.

35. Gupta M, Ravindranath S, Prasad D, Vidya TNC. Short-term variation in sex ratio estimates of Asian elephants due to space use differences between the sexes. Gajah 2016; 44: 5–15.

36. Kemf E, Santiapillai C. Asian elephants in the wild. Gland, Switzerland: WWF International’s Species Conservation Unit, WWF; 2000.

37. Blake S, Hedges S. Sinking the flagship: the case of forest elephants in Asia and Africa. Conserv Biol. 2004; 18: 1191–1202.

38. Schwarz CJ, Arnason AN. A general methodology for the analysis of capture-recapture experiments in open populations. Biometrics. 1996; 52: 860–873.

39. Pollock KH. A capture-recapture design robust to unequal probability of capture. J Wildl Manage. 1982; 46: 752–757.

40. Kendall WL, Pollock KH, Brownie C. A likelihood-based approach to capture-recapture estimation of demographic parameters under the robust design. Biometrics. 1995; 51: 293–308.

41. Kendall WL, Nichols JD, Hines JE. Estimating temporary emigration using capture-recapture data with Pollock's robust design. Ecology. 1997; 78: 563–578.

42. Norris III JL, Pollock KH. Nonparametric MLE under two closed capture-recapture models with heterogeneity. Biometrics. 1996; 52: 639–649.

43. Pledger S. Unified maximum likelihood estimates for closed capture-recapture models using mixtures. Biometrics. 2000; 56: 434–442.

44. Karanth KU, Nichols JD. Estimation of tiger densities in India using photographic captures and recaptures. Ecology. 1998; 79: 2852–2862.

45. Karanth KU, Nichols JD, Samba Kumar N. Estimating tiger abundance from camera trap data: field surveys and analytical issues. In: O'Connell AF, Nichols JD, Karanth KU, editors. Camera traps in animal ecology: methods and analyses. Tokyo: Springer; 2011. pp. 97–117.

46. Rees SG, Goodenough AE, Hart AG, Stafford R. Testing the effectiveness of capture mark recapture population estimation techniques using a computer simulation with known population size. Ecol Modell. 2011; 222: 3291–3294.

47. Otis DL, Burnham KP, White KP, Anderson DR. Statistical inference for capture data from closed populations. Wildlife Monogr. 1978; 62: 1–135.

48. Pollock KH, Nichols JD, Brownie C, Hines JE. Statistical inference for capture-recapture experiments. Wildlife Monogr. 1990; 107: 3–97.

49. Kendall WL. Robustness of closed capture-recapture methods to violations of the closure assumption. Ecology. 1999; 80: 2517–2525.

50. Harihar A, Pandav B, Goyal SP. Subsampling photographic capture-recapture data of tigers *(Panthera tigris)* to minimize closure violation and improve estimate precision: a case study. Popul Ecol. 2009; 51: 471–479.

51. Bondrup-Nielsen S. Density estimation as a function of live-trapping grid and home range size. Can J Zool. 1983; 61: 2361–2365.

52. Royle JA, Chandler RB, Sollmann R, Gardner B. Spatial capture-recapture. Waltham: Academic Press; 2014.

53. Sollmann R, Gardner B, Belant JL. How does spatial study design influence density estimates from spatial capture-recapture models? PLoS One. 2012; 7: e34575.

54. Calambokidis J, Barlow J. Abundance of blue and humpback whales in the eastern North Pacific estimated by capture-recapture and line-transect methods. Mar Mamm Sci. 2004; 20: 63–85.

55. Carroll EL, Patenaude N, Childerhouse SJ, Kraus SD, Fewster RM, Baker CS. Abundance of the New Zealand subantarctic southern right whale population estimated from photo-identification and genotype mark-recapture. Mar Biol. 2011; 158: 2565.

56. Smith HC, Pollock K, Waples K, Bradley S, Bejder L. Use of the robust design to estimate seasonal abundance and demographic parameters of a coastal bottlenose dolphin *(Tursiops aduncus)* population. PLoS One. 2013; 8: e76574.

57. Santostasi NL, Bonizzoni S, Bearzi G, Eddy L, Gimenez O. A robust design capture-recapture analysis of abundance, survival and temporary emigration of three Odontocete species in the Gulf of Corinth, Greece. PLoS One. 2016; 11: e0166650.

58. Bowden DC, Kufeld RC. Generalized mark-resight population size estimation applied to Colorado moose. J Wild Manage. 1995; 59: 840–851.

59. Weckerly FW. Constant proportionality in the female segment of a Roosevelt Elk population. J Wild Manage. 2007; 71:773–777.

60. MATLAB Release 2011a. Natick: The MathWorks, Inc; 2011.

61. Arivazhagan C, Sukumar R. Comparative demography of Asian elephant populations (*Elephas maximus*) in southern India. Centre for Ecological Sciences Technical Report No. 106. Bangalore: Centre for Ecological Sciences, Indian Institute of Science; 2005.

62. Vidya TNC, Prasad D, Ghosh A. Individual identification in Asian elephants. Gajah. 2014; 40: 3–17.

63. AERCC (Asian Elephant Research and Conservation Center). The Asian elephant in southern India: A GIS database for conservation of Project Elephant reserves. Technical Report No. 6. AERCC. Bangalore: Center for Ecological Sciences, Indian Institute of Science; 1998.

64. Baskaran N, Desai A. Ranging behaviour of the Asian elephant (*Elephas maximus*) in the Nilgiri Biosphere Reserve, South India. Gajah. 1996; 15: 41–57.

65. Baskaran N. Ranging and resource utilization by Asian elephant (*Elephas maximus* Linnaeus) in Nilgiri Biosphere Reserve, South India. Ph.D. Thesis, Bharathidasan University. 1998.

66. Payne KB, Langbauer Jr WR, Thomas EM. Infrasonic calls of the Asian elephant (*Elephas maximus*). Behav Ecol Sociobiol. 1986; 18: 297–301.

67. de Silva S, Webber CE, Weerathunga US, Pushpakumara TV, Weerakoon DK, et al. Demographic variables for wild Asian elephants using longitudinal observations. PloS One 2013; 8: e82788, doi:10.1371/journal.pone.0082788.

68. Laake JL. RMark: An R interface for analysis of capture-recapture data with MARK. AFSC Processed Rep 2013-01. Seattle: Alaska Fisheries Science Centre, NOAA, National Marine Fisheries Service; 2013. pp. 1–25.

69. White GC, Burnham KP. Program MARK: survival estimation from populations of marked animals. Bird Study. 1999; 46: S120–S139.

70. R Core Team. R: A language and environment for statistical computing. Vienna: R Foundation for Statistical Computing; 2014. Available: http://www.R-project.org/.

71. Ginsberg JR, Young TP. Measuring association between individuals or groups in behavioural studies. Anim Behav. 1992; 44: 377–379.

72. MacQueen J. Some methods for classification and analysis of multivariate observations. In: Le Cam LM, Neyman J, editors. Proceedings of the fifth Berkeley symposium on mathematical statistics and probability. Berkeley: University of California Press; 1967. pp. 281–297.

73. Seber GAF. Multivariate observations. New Jersey: John Wiley & Sons; 1984.

74. Rousseeuw PJ. Silhouettes: a graphical aid to the interpretation and validation of cluster analysis. J Comput Appl Math. 1987; 20: 53–65.

75. StatSoft, Inc. STATISTICA for Windows [computer program manual]. Tulsa: StatSoft, Inc; 1996.

